# Connecting omics signatures of diseases, drugs, and mechanisms of actions with iLINCS

**DOI:** 10.1101/826271

**Authors:** Marcin Pilarczyk, Michal Kouril, Behrouz Shamsaei, Juozas Vasiliauskas, Wen Niu, Naim Mahi, Lixia Zhang, Nicholas Clark, Yan Ren, Shana White, Rashid Karim, Huan Xu, Jacek Biesiada, Mark F. Bennett, Sarah Davidson, John F Reichard, Kurt Roberts, Vasileios Stathias, Amar Koleti, Dusica Vidovic, Daniel J.B. Clarke, Stephan C. Schurer, Avi Ma’ayan, Jarek Meller, Mario Medvedovic

## Abstract

There are only a few platforms that integrate multiple omics data types, bioinformatics tools, and interfaces for integrative analyses and visualization that do not require programming skills. Among these, iLINCS is unique in scope and versatility of the data provided and the analytics facilitated. iLINCS (http://ilincs.org) is an integrative web-based platform for analysis of omics data and signatures of cellular perturbations. The platform facilitates analysis of user-submitted omics signatures of diseases and cellular perturbations in the context of a large compendium of pre-computed signatures (>200,000), as well as mining and re-analysis of the large collection of omics datasets (>12,000), pre-computed signatures, and their connections. Analytics workflows driven by user-friendly interfaces enable users with only conceptual understanding of the analysis strategy to execute sophisticated analyses of omics signatures, such as systems biology analyses and interpretation of signatures, mechanism of action analysis, and signature-driven drug repositioning. In summary, iLINCS workflows integrate vast omics data resources and a range of analytics and interactive visualization tools into a comprehensive platform for analysis of omics signatures.

## Background

Transcriptomics and proteomics (omics) signatures in response to cellular perturbations consist of changes in gene or protein expression levels after the perturbation. An omics signature is a high-dimensional readout of cellular state change that provides information about the biological processes affected by the perturbation lead to the post-perturbation phenotype of the cell. The signature on its own provides information, although not always directly discernable, about the molecular mechanisms by which the perturbation causes observed changes. If we consider a disease to be a perturbation of the homeostatic biological system under normal physiology, then the omics signature of a disease are the differences in gene/protein expression levels between disease and non-diseased tissue samples.

The low cost and effectiveness of transcriptomics assays^1-4^ has resulted in an abundance of transcriptomics datasets and signatures. Recent advances in the field of high throughput proteomics made generation of large numbers of proteomics signatures a reality^5,6^. Several recent efforts were directed at systematic generation of omics signatures of cellular perturbations^7^ and at generating libraries of signatures by re-analyzing public domain omics datasets^8,9^. The recently released library of integrated network-based cellular signatures (LINCS)^7^ L1000 dataset generated transcriptomic signatures at unprecedented scale^2^. The availability of resulting libraries of signatures open exciting new avenues for learning about the mechanisms of diseases and the search for effective therapeutics^10^.

The analysis and interpretation of omics signatures has been intensely researched. Numerous methods and tools have been developed for identifying changes in molecular phenotypes implicated by transcriptional signatures based on gene set enrichment, pathway, and network analyses approaches^11-13^. Directly matching transcriptional signatures of a disease with negatively correlated transcriptional signatures of chemical perturbations (CP) underlies the *Connectivity Map* (CMap) approach to identifying potential drug candidates^10,14,15^. Similarly, correlating signatures of chemical perturbagens with genetic perturbations of specific genes has been used to identify putative targets of drugs and chemical perturbagens ^2^.

To fully exploit the information contained within omics signature libraries and within countless omics signatures generated every day by investigators around the world, new user-friendly integrative tools are needed that bring this data together, and are accessible to a large segment of biomedical research community. The integrative LINCS (iLINCS) portal brings together libraries of precomputed signatures, formatted datasets, connections between signatures, and integrates them with a bioinformatics analysis engine into a coherent system for omics signature analysis.

## Results

iLINCS (available http://ilincs.org) is an integrative user-friendly web platform for the analysis of omics (transcriptomic and proteomic) datasets and signatures of cellular perturbations. The key components of iLINCS are: *Interactive and interconnected analytical workflows for creation and analysis of omics signatures; The large collection of datasets*, pre-computed *signatures* and their *connections; and user-friendly graphical interfaces for executing analytical tasks and workflows*. The central concept in iLINCS is the omics signature which can be retrieved from the pre-computed signature libraries within the iLINCS database, submitted by the user, or constructed using one of the iLINCS datasets (Fig 1A). The signatures in iLINCS consist of differential gene or protein expression levels and associated p-values between perturbed and baseline samples for all, or any subset of measured genes/proteins. Signatures submitted by the user can also be in the form of a list of genes/proteins, or a list of up- and down-regulated genes/proteins. Analytical workflows facilitate systems biology interpretation of the signature (Fig 1B) and the connectivity analysis of the signature with all iLINCS pre-computed signatures (Fig 1C). Connected signatures can further be analyzed in terms of the patterns of gene/protein expression level changes that underlie the connectivity with the query signature, or through the analysis of gene/protein targets of connected perturbagens (Fig 1D). Ultimately, the multi-layered systems biology analyses, and the connectivity analyses lead to biological insights, and identification of therapeutic targets and putative therapeutic agents (Fig 1E). Below we provide an overview of the key data and analytic components of iLINCS, and then we present three use cases to demonstrate iLINCS’ capacity to generate impactful results.

**Fig 1.**
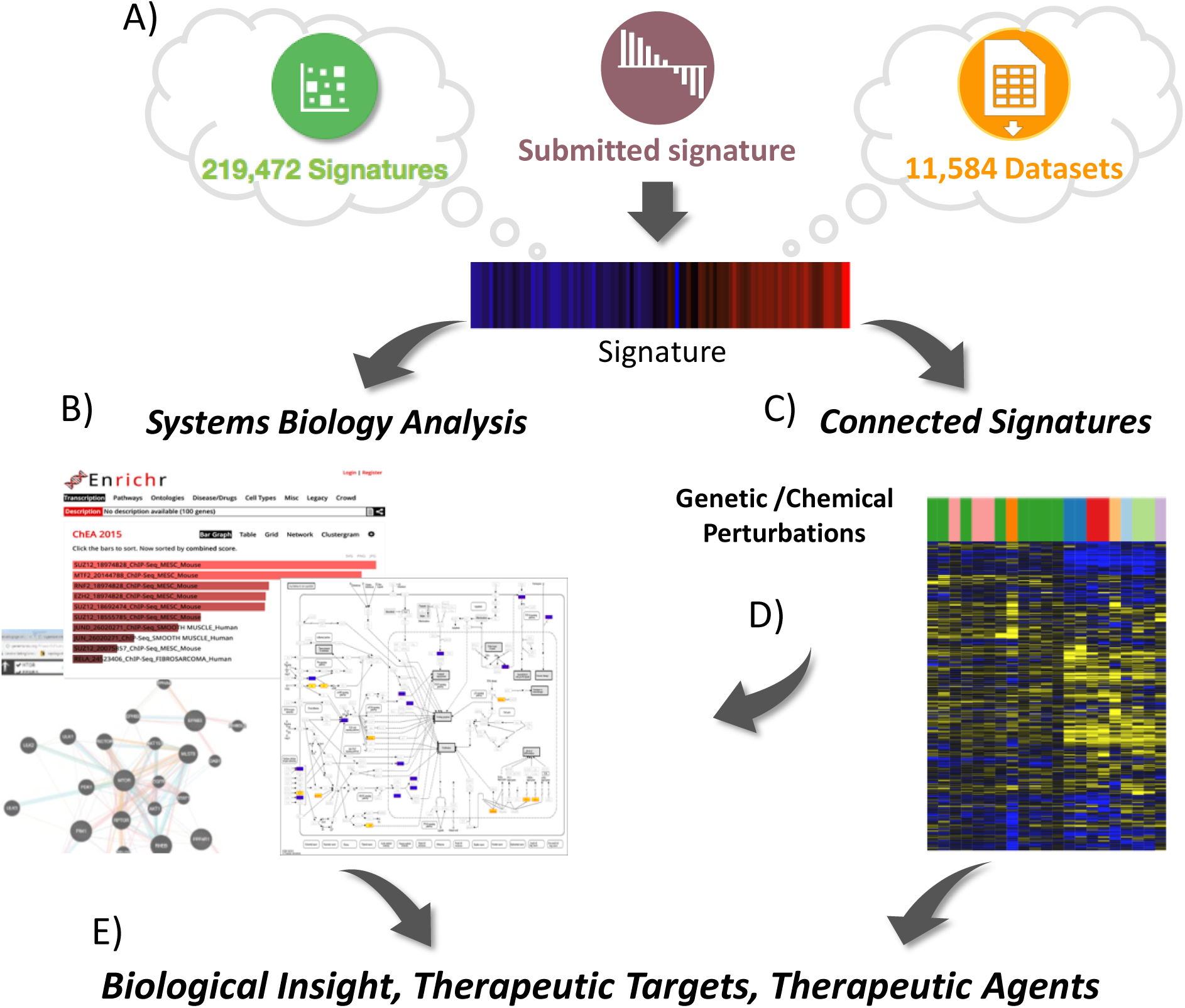
Integrative omics signature analysis in iLINCS. A) A signature can be selected by querying the iILINCS database, submitted by the user, or constructed by analyzing an iLINCS omics dataset. B) The signature can be analyzed using a range of systems biology methods (gene set enrichment, pathway and network analyses). C) Signature “connectivity” analyses can be applied to identify cellular perturbations and biological states of similar signatures. D) The analysis of connected signatures, as well as the identity of the perturbed genes and proteins leading to the connected signatures, can be used to elucidate mechanisms of action. E) Ultimately, the results of the analyses lead to insights and hypotheses about potential therapeutic targets and therapeutic agents.

### Interconnected workflows for constructing and analyzing omics signatures

Interactive analytical workflows in iLINCS facilitate signature construction through differential expression analysis as well as clustering, dimensionality reduction, functional enrichment, signature connectivity analysis, pathway and network analysis, and integrative interactive visualization. Visualizations include interactive scatter plots, volcano and GSEA plots, heatmaps, and pathway and network node and stick diagram (Supplemental Figure 1). Users can download raw data and signatures, analysis results and publication-ready graphics. iLINCS internal analysis and visualization engine uses R ^16^, Bioconductor packages^17^, the Shiny framework^18^, interactive graphics created with *ggplot* ^19^and *plotly* ^*20*^, and integration of open-source visualization tools such as FTreeView^21^ and Morpheous ^22^. iLINCS also facilitates seamless integration with a wide range of task-specific online bioinformatics and systems biology tools and resources including Enrichr^23^, DAVID^24^, ToppGene^25^, Reactome^26^, KEGG^27^, GeneMania^28^, X2K Web^29^, L1000FWD^30^, STITCH^31^, Clustergrammer^32^, piNET ^33^, LINCS Data Portal^34^, ScrubChem^35^, PubChem^36^, GEO^37^, ArrayExpress^38^ and GREIN^39^. Programmatic access to iLINCS data, workflows and visualizations are facilitated by embedding the calls to iLINCS API which are documented with the Swagger community standard. Examples of utilizing the iLINCS API within data analysis scripts are provided on GitHub (https://github.com/uc-bd2k/ilincsAPI). The iLINCS software architecture is described in Supplemental Figure 2.

**Fig 2.**
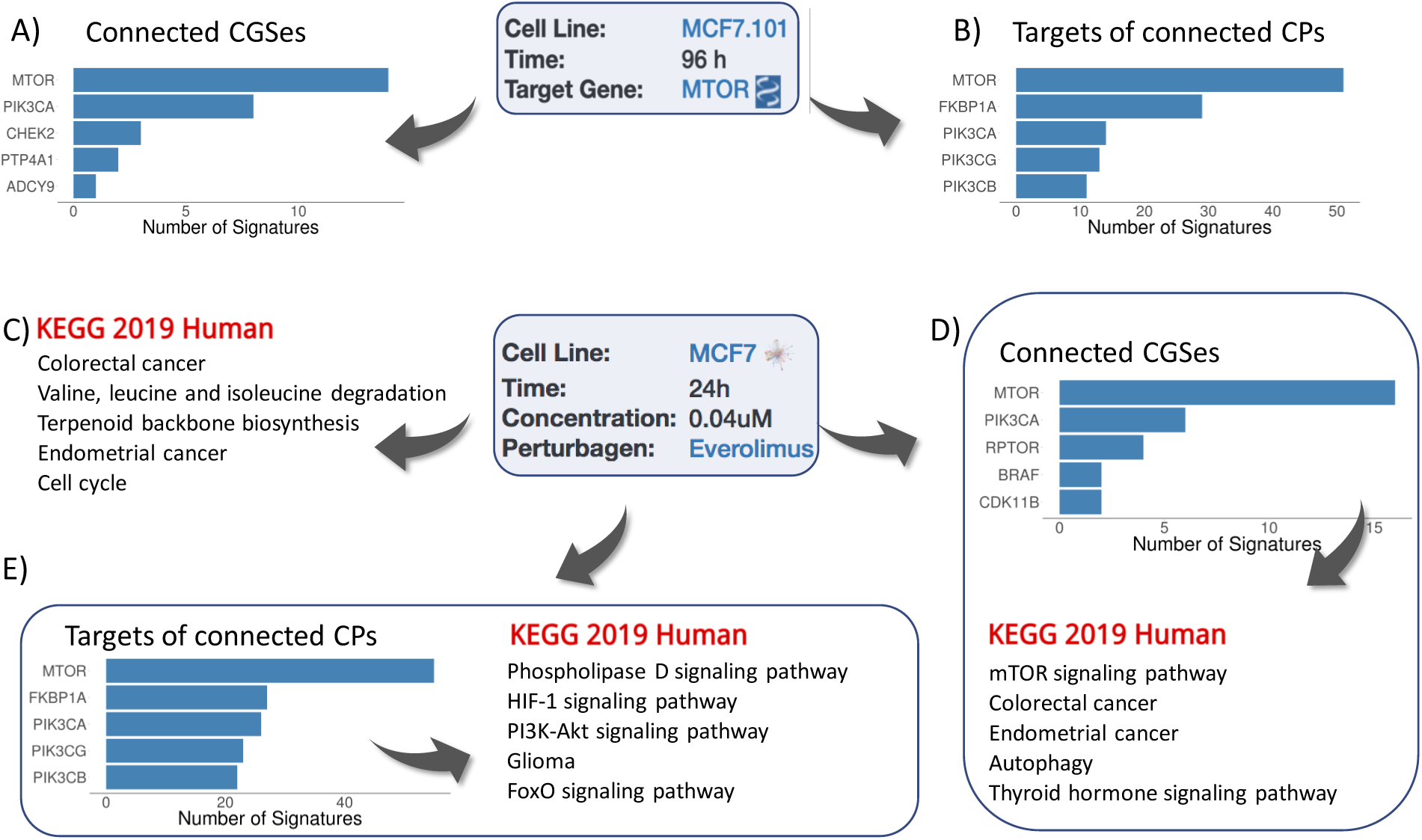
Analysis of LINCS L1000 signatures of genetic and chemical perturbations. A) Most frequently perturbed genes among the Consensus Genes Signatures (CGS) connected to the mTOR knock-down CGS; B) Most frequent inhibition targets of chemical perturbagens with signatures connected to the mTOR CGS signature; C) Most enriched biological pathways for the everolimus signature; D) Most frequently perturbed genes among CGSes connected with everolimus signature, and pathways most enriched by the perturbed genes; E) Most frequent inhibition targets of chemical perturbagens with signatures connected to the everolimus signature and the pathways most enriched by the genes of the targeted proteins.

### iLINCS libraries of datasets, signatures and connections

iLINCS backend ***Databases*** contain >10,000 processed omics datasets, >220,000 omics signatures and >10^9^ statistically significant “connections” between signatures. ***Omics datasets*** available for analysis and signatures creation include transcriptomic (RNA-seq and microarray) and proteomic (Reverse Phase Protein Arrays ^40^ and LINCS targeted mass spectrometry proteomics^5^) datasets. Dataset collections include transcriptomic and proteomics data generated by The Cancer Genome Atlas (TCGA) project, GEO GDS datasets, and the complete collection of GEO RNA-seq datasets. ***Omics signatures include***: *LINCS chemical and genetic perturbation signatures* consisting of genome-wide transcriptional response after genetic loss of function perturbation of more than 3,500 genes, or a perturbation by one of more than 4,000 chemical perturbagens based on LINCS L1000 assay data ^2^, *DrugMatrix* Chemogenomic database of 5,200 transcriptomic profiles of chemical toxicity ^41^, *Disease Related Signatures* consisting of 9,000 transcriptional signatures constructed by comparing sample groups within the collection of curated transcriptomics datasets from GEO^42^, EBI Expression Atlas^8^ signatures, and 5,000 pharmacogenomics signatures constructed from public domain datasets^4,43^.

### Use case 1: Identifying chemical perturbagens emulating genetic perturbation of the mTOR gene

Aberrant activation of mTOR signaling underlies multiple human diseases and numerous efforts in designing drugs that modulate activity of mTOR signaling are under way^44^. Here we use the signature of genetic perturbation (via CRISPR knock-down) of the mTOR genes to identify chemical perturbagens mimicking the perturbation of the mTOR genes. First, we search through iLINCS libraries for Consensus Genes Signatures (CGSes) of mTOR knock-down and use the CRISPR CGS in MCF-7 cell line as the query signature. The connectivity analysis identifies 258 LINCS CGSes and 831 CP Signatures with statistically significant correlation with the query signature. Top 100 most connected CGSes are dominated by the signatures of genetic perturbations of mTOR and PIK3CA genes (Fig 2A), whereas all top 5 most frequent inhibition targets of CPs among top 100 most connected CP signatures are mTOR and PIK3 proteins (Fig 2B). Results clearly indicate that the query mTOR CGS is highly specific and sensitive to perturbation of the mTOR pathway and effectively identifies chemical perturbagens capable of inhibiting mTOR signaling. The full list of connected signatures is shown in Supplemental Table ST1. The connected CP signatures also include several chemical perturbagens with highly connected signatures that have not been known to target mTOR signaling providing additional candidate inhibitors. Step by step instructions for performing this analysis in iLINCS are provided in Supplemental Workflow SW1.

### Use case 2: Mechanism of action analysis via connection to genetic perturbation signatures

Identifying small molecules (i.e. chemical perturbagens) that can modulate activity of disease-related proteins or pathways is the cornerstone of intelligent drug design. Transcriptional signature of chemical perturbagens often carry only an echo of such effects since the proteins directly targeted by a chemical and the interacting signaling proteins are not transcriptionally changed. iLINCS offers the solution for this problem by connecting the CP signatures to LINCS CGSes, and facilitating a follow-up systems biology analysis of genes whose CGSes are highly correlated with the CP signature. This strategy is illustrated by the analysis of the signature of the 24 hour, 0.04µM treatment of the MCF-7 cell line with the mTOR inhibitor everolimus (Fig 2CDE). Step by step instructions for performing this analysis in iLINCS are provided in Supplemental Workflow SW2. Traditional pathway enrichment analysis of this CP signature via iLINCS connection to Enrichr (Fig 2C) fails to identify the mTOR pathway as being affected. In the next step, we first connect the CP signature to LINCS CGSes and then perform pathway enrichment analysis of genes with correlated CGSes. This analysis correctly identifies mTOR signaling pathway as the top affected pathway (Fig 2D). Similarly, connectivity analysis with other CP signatures followed by the enrichment analysis of protein targets of top 100 most connected CPs again identifies the Pi3k-Akt signaling pathway as one of the most enriched (Fig 2E). In conclusion, both pathway analysis of differentially expressed genes in the everolimus signature and pathway analysis of connected genetic and chemical perturbagens provide us with important information about effects of everolimus. However, only the analyses of connected perturbagens correctly pinpoints the direct mechanism of action of the everolimus, which is the inhibition of mTOR signaling.

### Use case 3: Proteo-genomics analysis of cancer driver events in breast cancer

We analyzed TCGA breast cancer RNA-seq and RPPA data using the iLINCS “Datasets” workflow to construct the differential gene and protein expression signatures contrasting Luminal A and Her2 enriched (Her2E) breast tumors^45^. The protein expression signature immediately implicated known driver events distinguishing the two subtypes, the Luminal A cancers being driven by abnormal activity of the estrogen receptor and the Her2E tumors driven by abnormally high activity of the Her2 protein (Fig 3A). In addition to expected proteins, the increased level of phosphorylated EIF4EBP1 protein may indicate increased level of mTOR signaling in Her2E tumors.

**Fig 3.**
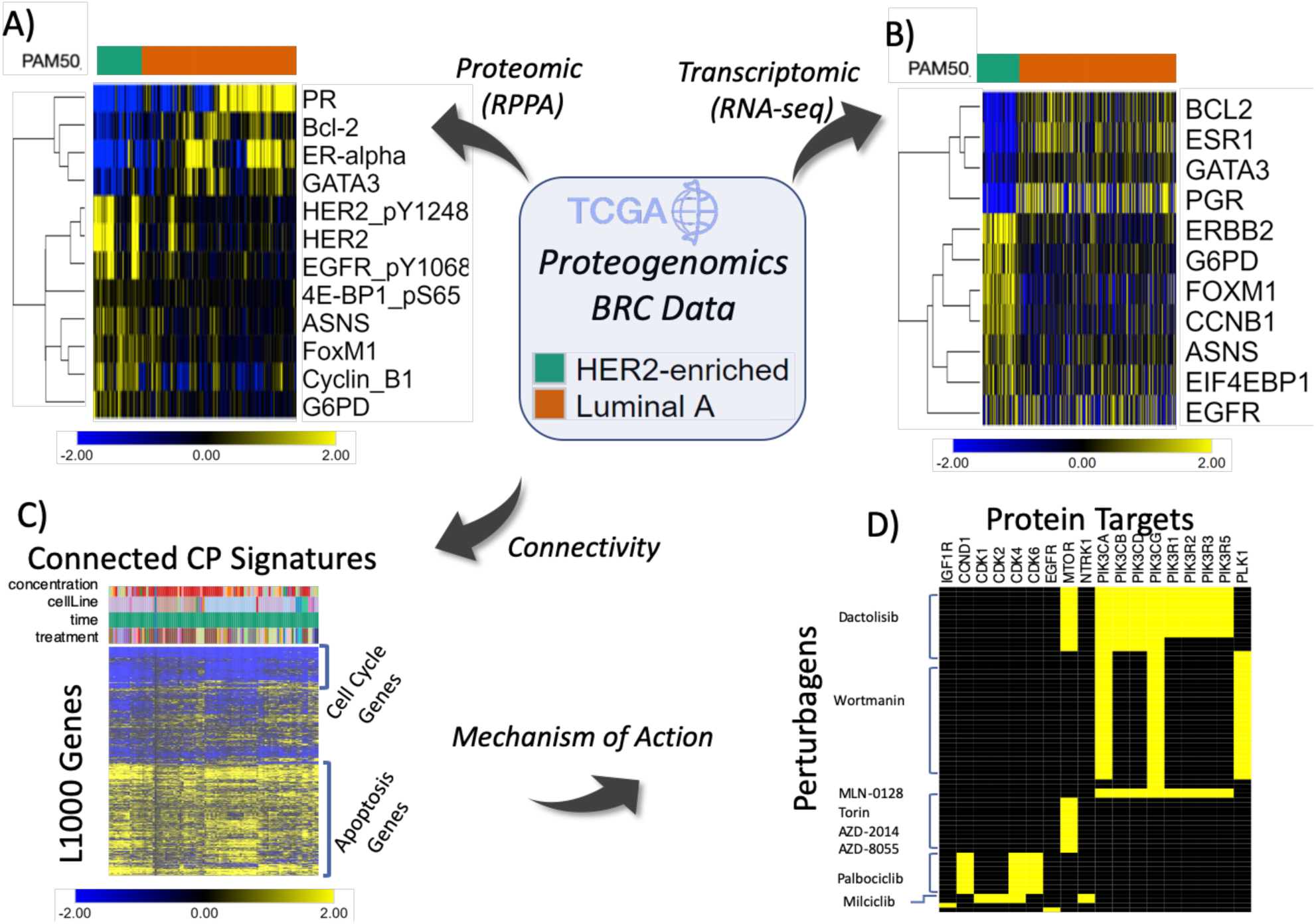
Proteo-genomics analysis of cancer driver events in breast cancer. A) Most differentially expressed proteins in the proteomics signatures constructed by comparing RPPA profiles of Her2E and Luminal A BRC samples; B) Gene expression profile of the genes corresponding to proteins in A) based on RNA-seq data; C) Top 100 CP signatures most connected with the transcriptional signature constructed by comparing RNA-seq profiles of Her2E and Luminal A samples; D) Selected chemical perturbagens and their targets for CP signatures in C).

The corresponding RNA-seq signature showed similar patterns of expression of key genes (Fig 3B). All genes were differentially expressed (Bonferroni adjusted p-value<0.01) except for EGFR, indicating that the difference in levels of post-translationally activated (phosphorylated) versions of the proteins may have come from activation of upstream kinases instead of overall increase in gene/protein expression. Analysis of 665 most significantly upregulated genes in Her2E tumors (p-value<1e-10) identified cell cycle-related KEGG pathways to be the most significantly(Cell cycle, p-value=1.3e-26; DNA replication, p-value=1.4e-13) enriched according to the Enrichr combined scores (See Supplemental Table ST2 for all results). This implicates a known increased proliferation of Her2E tumors in comparison to Luminal A tumors^46^. The connectivity analysis of the RNA-seq signature with LINCS CP signatures shows that treating several different cancer cell lines with inhibitors of PI3K, mTOR, CDK (Fig 3C) and inhibitors of some other more generic proliferation targets (eg. TOP21, AURKA) (see Supplemental Table ST3 for complete results) produces signatures that are positively correlated with RNA-seq Luminal A vs Her2E signature, indicating that such treatments are pushing cancer cell lines toward greater phenotypic similarity with Luminal A tumors.

Group analysis of connected CP signatures (Fig 3C) indicates that results of connectivity analysis may be largely driven by the inhibition of proliferation as evidenced by enrichment of down-regulated genes by cell cycle genes and up-regulated genes by apoptosis related genes in connected signatures. However, the dominance of PI3K and mTOR inhibitor signatures (Fig 3D) suggests that the connections, to some extent, may also be driven by more specific targeting of PI3K-mTOR signaling, which may be more active in Her2E cancers as indicated by increased levels of the phosphorylated EIF4EBP1 protein.

High positive connectivity to Cancer Therapeutic Response Signatures^47^ of two PI3K inhibitors in breast cancer cell lines also indicates that Her2E cancers may be more sensitive to mTOR inhibition (Supplemental Table ST3). Connectivity analysis with ENCODE transcription factor targets signatures recapitulated known biology (negative association with E2F4 binding signatures implicating higher proliferation of Her2E tumors and positive associations with ER*α* binding signatures, implicating the increase ER*α* activity in Luminal A tumors) (Supplemental Table ST3). Most connected signatures in analysis of Disease related signatures^42^ extracted from GEO data and EBI Expression Atlas^8^ signatures were all related to comparisons of different breast cancers samples (Supplemental Table ST3). Step by step instructions for performing this analysis in iLINCS are provided in Supplemental Workflow SW3.

#### Other use cases

The three interconnected iLINCS workflows (Signatures, Datasets, Genes), facilitate a wide range of possible use cases. The three detailed cases above all use either pre-computed iLINCS signatures, or iLINCS omics datasets to construct signatures. Querying iLINCS with user submitted external signatures, genes and gene lists allows identification of connected perturbations and signatures. It also allows users to answer more specific questions about expression patterns of genes or gene lists of interest in specific datasets or across a class of cellular perturbations. For example, a query with a specific gene of interest can identify sets of perturbations that significantly affect the expression of the gene and thus offering a set of chemicals, or genetic perturbations that can be used to modulate the activity of the corresponding protein. A query with a list of genes whose coordinated expression is known to be a hallmark of a specific biological state or process^48^ can identify a set of perturbations that can accordingly modify cell phenotype. Additional use cases have also been illustrated in several published scientific studies utilizing iLINCS: identification of putative therapeutic agents for schizophrenia^49,50^, developing new strategies for ERα degradation in breast cancers^51^, inhibiting the protective effects of stromal cells against chemotherapy in breast cancer^52^ and rational drug combination design to inhibit epithelial-mesenchymal transition^53^.

## Discussion

iLINCS is a unique integrated platform for analysis of omics signatures. The three use cases described here only scratch the surface of the wide range of possible analyses facilitated by the interconnected analytical workflows and the large collections of omics datasets, signatures and their connections. These cases also feature only a subset of all analytical tools integrated within the iLINCS platform. The user interfaces are streamlined and strive to be self-explanatory to most scientists with conceptual understanding of omics data analysis. All analyses presented here were performed by typing the initial queries and then using the mouse to navigate user interfaces without ever having to copy and/or re-submit any portions of the data and results to a separate analytical tool. iLINCS implements the complete systematic polypharmacology and drug repurposing ^54^ workflow, and provides new innovative workflows for harnessing the full potential of LINCS omics signatures.

In addition to facilitating standard analyses, iLINCS also implements innovative workflows for biological interpretation of omics signatures via connectivity analysis. For example, in use case 2 we show how connectivity analysis coupled with pathway and gene set enrichment analysis can implicate mechanism of action of a chemical perturbagen when standard enrichment analysis applied to the differentially expressed genes fails to recover targeted signaling pathways. In a similar vein, iLINCS has been successfully used to identify putative therapeutic agents by connecting changes in proteomics profiles in neurons from patients with schizophrenia; first with the LINCS CGSes of the corresponding genes, and then with LINCS CP signatures^50^. These analyses led to identification of PPAR agonists as promising therapeutic agents capable of reversing bioenergetic signature of schizophrenia, which were subsequently shown to modulate behavioral phenotypes in rat model of schizophrenia^49^. In last several months, iLINCS has also been used to research immune response to COVID-19 infection and new therapeutic approaches ^55 56 57 58 59^.

Several online tools have been developed for the analysis and mining LINCS L1000 signature libraries. They facilitate online queries of L1000 signatures^60,61^ and construction of scripted pipelines for in-depth analysis of transcriptomics data and signatures^62^. The LINCS Transcriptomic Center at the Broad Institute developed the *clue*.*io* query tool deployed by the Broad Connectivity Map team which facilitates connectivity analysis of user submitted signatures^2^. iLINCS replicates the connectivity analysis functionality, and indeed, the equivalent queries of the two systems may return qualitatively similar results (see Supplemental Results for a use case comparison). However, the scope of iLINCS is much broader. It provides connectivity analysis with signatures beyond Connectivity Map datasets and provides many primary omics datasets for users to construct their own signatures. Furthermore, analytical workflows in iLINCS facilitate deep systems biology analysis and knowledge discovery of both omics signatures and the genes and protein targets identified through connectivity analysis. Comparison to several other web resources that partially cover different aspects of *iLINCS* functionality are summarized in Supplemental Table S4.

iLINCS removes technical roadblocks for users without programming background to re-use a large fraction of publicly available omics datasets and signatures. Recent effort in terms of standardizing ^63^ and indexing ^64^ efforts are improving findability and re-usability of public domain omics data. iLINCS is taking the next logical step in integrating public domain data and signatures with user-friendly analysis toolbox. Furthermore, all analyses steps behind the iLINCS UI’s are driven by API which themselves can and have been already used within computational pipelines based on scripting languages^65^, such as R, Python and JavaScript, or to power functionality of other web analysis tools^33,66^. This makes iLINCS a natural tool for analysis and interpretation of omics signatures for scientists preferring point-and-click GUIs as well as data scientists using scripted analytical pipelines.

## Methods

### Perturbation signatures

All pre-computed perturbation signatures in iLINCS, as well as signatures created using an iLINCS dataset, consist of two vectors: the vector of log-scale differential expressions between the perturbed samples and baseline samples ***d***=(*d1,…,dN*), and the vector of associated p-values ***p***=(*p1*,…,*pN*), where N is the number of genes or proteins in the signature. Signatures submitted by the user can also consist of only log-scale differential expressions without p-values, list of up-and down-regulated genes, and single list of genes.

### Signature connectivity analysis

Depending on the exact type of the query signature, the connectivity analysis with libraries of pre-computed iLINCS signatures are computed using different connectivity metric. The choice of the similarity metric to be used in different contexts was driven by benchmarking six different methods (Supplementary Result 2).

If the query signature is selected from iLINCS libraries of pre-computed signatures, the connectivity with all other iLINCS signatures is pre-computed using the extreme Pearson’s correlation^67,68^ of signed significances of all genes. The signed significance of the i^th^ gene is defined as

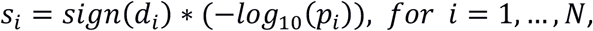

and the signed significance signature is ***s***=(*s1,…,sN*). The extreme signed signature ***e***=(*e1,…,eN*) is then constructing by setting the signed significances of all genes other than the top 100 and bottom 100 to zero:

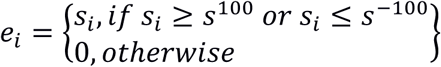

Where *s*^100^ is the 100^th^ most positive *si* and *s*^-100^ is the 100^th^ most negative *si*. The extreme Pearson correlation between two signatures is then calculated as the standard Pearson’s correlation between the extreme signed significance signatures.

If the query signature is created from an iLINCS dataset, or directly uploaded by the user, the connectivity with all iLINCS signatures is calculated as the weighted correlation between the two vectors of log-differential expressions and the vector of weights equal to [-log10(p-value of the query) - log10(p-value of the iLINCS signature)]^69^. When the user-uploaded signature consists of only log differential expression levels without p-values, the weight for the correlation is based only on the p-values of the iLINCS signatures [-log10(p-values of the iLINCS signatures)].

If the query signature uploaded by the user consists of the lists of up- and down-regulated genes connectivity is calculated by assigning -1 to down-regulated and +1 upregulated genes and calculating Pearson’s correlation between such vector and iLINCS signatures. The calculated statistical significance of the correlation in this case is equivalent to the t-test for the difference between differential expression measures of iLINCS signatures between up- and down-regulated genes.

If the query signature is uploaded by the user in a form of a gene list, the connectivity with iLINCS signatures is calculated as the enrichment of highly significant differential expression levels in iLINCS signature within the submitted gene list using the Random Set analysis ^70^.

### Perturbagen connectivity analysis

The connectivity between a query signature and a “perturbagen” is established using the enrichment analysis of individual connectivity scores between the query signature and set of all L1000 signatures of the perturbagen (for all cell lines, time points and concentrations). The analysis establishes whether the connectivity scores as a set are “unusually” high based on the Random Set analysis^70^.

### iLINCS signature libraries

#### LINCS L1000 signature libraries (Consensus gene knockdown signatures (CGS), Overexpression gene signatures and Chemical perturbation signatures)

For all LINCS L1000 signature libraries, the signatures are constructed by combining the Level 4, population control signature replicates from two released GEO datasets (GSE92742 and GSE70138) into the Level 5 moderated Z scores (MODZ) by calculating weighted averages as described in the primary publication for the L1000 Connectivity Map dataset^2^. Only signatures showing evidence of being reproducible by having the 75th quantile of pairwise spearman correlations of level 4 replicates (Broad institute distil_cc_q75 quality control metric^2^) greater than 0.2 are included. The corresponding p-values were calculated by comparing MODZ of each gene to zero using the Empirical Bayes weighted t-test with the same weights used for calculating MODZs. The shRNA and CRISPR knock-down signatures targeting the same gene were further aggregated into Consensus gene signatures (CGSes)^2^ by the same procedure used to calculate MODZs and associated p-values.

#### LINCS targeted proteomics signatures

Signatures of chemical perturbations assayed by the quantitative targeted mass spectrometry proteomics P100 assay measuring levels 96 phosphopeptides and GCP assay against ∼60 probes that monitor combinations of post-translational modifications on histones^5^.

#### Disease related signatures

Transcriptional signatures constructed by comparing sample groups within the collection of curated public domain transcriptional dataset (GEO DataSets collection)^37^. Each signature consists of differential expressions and associated p-values for all genes calculated using Empirical Bayes linear model implemented in the *limma* package.

#### ENCODE transcription factor binding signatures

Genome-wide transcription factor (TF) binding signatures constructed by applying the TREG methodology to ENCODE ChiP-seq^71^. Each signature consists of scores and probabilities of regulation by the given TF in the specific context (cell line and treatment) for each gene in the genome.

#### Connectivity Map Signatures

Transcriptional signatures of perturbagen activity constructed based on the version 2 of the original Connectivity Map dataset using Affymetrix expression arrays^42^. Each signature consists of differential expressions and associated p-values for all genes when comparing perturbagen treated cell lines with appropriate controls.

#### DrugMatrix signatures

Toxicogenomic signatures of over 600 different compounds^41^ maintained by the National Toxicology Program^72^ consisting of genome-wide differential gene expression levels and associated p-values.

#### Transcriptional signatures from EBI Expression Atlas

All mouse, rat and human differential expression signatures and associated p-values from manually curated comparisons in the Expression Atlas^8^.

#### Cancer therapeutics response signatures

These signatures were created by combining transcriptional data with drug sensitivity data from the Cancer Therapeutics Response Portal (CTRP) project^47^. Signatures were created separately for each tissue/cell lineage in the dataset by comparing gene expression between the five cell lines of that lineage that were most and five that were least sensitive to a given drug area as measured by the concentration-response curve (AUC) using two-sample t-test.

#### Pharmacogenomics transcriptional signatures

These signatures were created by calculating differential gene expression levels and associated p-value between cell-lines treated with anti-cancer drugs and the corresponding controls in two separate projects: The NCI Transcriptional Pharmacodynamics Workbench (NCI-TPW)^43^ and the Plate-seq project dataset^4^.

### Constructing signatures from iLINCS datasets

The transcriptomics or proteomics signature is constructed by comparing expression levels of two groups of samples (treatment group and baseline group) using Empirical Bayes linear model implemented in the *limma* package^73^. For the *GREIN* collection of GEO RNA-seq datasets^74^, the signatures are constructed using the negative-binomial generalized linear model as implemented in the *edgeR* package^75^.

### Analytical tools, web applications and web resources

Signatures analytics in iLINCS is facilitated via native R, Java, JavaScript and Shiny applications, and via API connections to external web application and services. Brief listing of analysis and visualization tools is provided here. The overall structure of iLINCS is described in the Supplemental Results.

*Gene list enrichment analysis* is facilitated by directly submitting lists of gene to any of the three prominent enrichment analysis web tools: Enrichr^23^, DAVID^24^, ToppGene^25^. The manipulation and selection of list of signature genes is facilitated via an interactive volcano plot JavaScript application (shown in Supplemental Workflow 3).

*Pathway analysis* is facilitated through either general purpose enrichment tool (Enrichr, DAVID, ToppGene), the enrichment analysis of Reactome pathways via Reactome online tool^26^, and internal R routines for SPIA analysis^76^ of KEGG pathways and general visualization of signatures in the context of KEGG pathways using the KEGG API^27^.

*Network analysis* is facilitated by submitting lists of genes to Genemania^28^ and by internal iLINCS Shiny Signature Network Analysis (SigNetA) application.

*Heatamap visualizations* are facilitated by native iLINCS applications: Java based FTreeView^21^, modified version of the JavaScript based Morpheus^22^ and a Shiny based HeatMap application and by connection to the web application Clustergrammer^32^.

*Dimensionality reduction analysis (PCA and t-SNE*^*77*^) and visualization of high-dimensional relationship via interactive 2D and 3D scatter plots is facilitated via internal iLINCS Shiny applications.

*Interactive box-plots, scatter plots, GSEA plots, bar charts and pie charts* used throughout iLINCS are implemented using R ggplot^19^ and plotly^20^.

*Additional analysis are provided by connection* X2K Web^29^ (inference of upstream regulatory networks from signature genes), L1000FWD^30^ (connectivity with signatures constructed using characteristic dimension methodology), STITCH^31^ (visualization of drug-target networks), piNET^33^ (visualization of gene-to-pathway relationships for signature genes).

*Additional information about drugs, genes and proteins* are provided by links to, LINCS Data Portal^78^, ScrubChem^35^, PubChem^36^, Harmonizome^79^, GeneCards^80^, and several other only databases.

### Gene and protein expression dataset collections

iLINCS backend databases provide access to more than 11,000 pre-processed gene and protein expression datasets that can be used to create and analyze gene and expression protein signatures. Datasets are thematically organized into eight collections with some datasets assigned to multiple collections. User can search all datasets or browse datasets by collection.

#### LINCS collection

Datasets generated by the LINCS data and signature generation centers^7^

#### TCGA collection

Gene expression (RNASeqV2), protein expression (RPPA), and copy number variation data generated by TCGA project^45^

#### GDS collection

A curated collection of GEO Gene Datasets (GDS)^37^

#### Cancer collection

An ad-hoc collection of cancer related genomics and proteomic datasets

#### Toxicogenomics collection

An ad-hoc collection of toxicogenomics datasets

#### RPPA collection

An ad-hoc collection of proteomic datasets generated by Reverse Phase Protein Array assay^81^

#### GREIN collection

Complete collection of preprocessed human, mouse and rat RNA-seq data in GEO provided by the GEO RNA-seq Experiments Interactive Navigator (GREIN)^74^

#### Reference collection

An ad-hoc collection of important gene expression datasets.

## Acknowledgements

We would like to acknowledge the big contribution by Dr. Mehdi Fazel Najafabadi in the development of the backend R-based analytical engine. Routines developed by Dr. Najafabadi perform most of the statistical analyses and create a portion of interactive visualizations displayed in iLINCS. We would also like to acknowledge contributions of Dr. Vineet Joshi and Dr. Mukta Phatak for development of early prototypes of the iLINCS portal. The development of iLINCS was funded by the LINCS-BD2K Data Coordination and Integration Center U54 grant (U54HL127624), Center for Environmental Genetics P30 grant (P30ES006096) and the Computational Tool Development and Integrative Data Analysis for LINCS U01 grant (U01HL111638).

