## Supplementary materials for "Connecting omics signatures of diseases, drugs, and mechanisms of actions with iLINCS"

### Supplemental Figure 1: Interactive visualization tools in iLINCS.

A) Interactive heatmaps: As a gold standard graphical display for visualizing high-dimensional data and relationships, interactive heatmaps are used throughout iLINCS via several different applications: Native Shiny heatmap, Java based FTreeView, Java script based Morpheus and Clustergrammer. Diversity of heatmap apps provide for diversity of functionalities and it facilitates use of iLINCS in different configurations of network speed vs the computer speed. B) Interactive volcano plots are used for visualizing and selecting informative (eg differentially expressed) genes/proteins in a signature; C) Interactive scatter plots are used to visualize relationships between two signatures and identifying genes/proteins driving the "connectivity"; D) Interactive box plots are used to visualize differential distribution of gene/proteins in different samples and up- and down-regulated genes/proteins in different signatures; E) Interactive GSEA plots serve to visualize strength of connectivity and identify genes/proteins driving the connectivity between a gene list and a signature; F) Interactive 3D scatter plots are used to visualize high-dimensional relationship in dimensionality reduction analysis (PCA and t-SNE); G) Connected 2D scatter plots for visualizing high-dimensional relationship in dimensionality reduction analysis (PCA and t-SNE); H) Interactive pathway visualizations allows for exploring the relationships between signatures and pathways; and K) Interactive network visualizations is used to integrate signatures with the global protein-protein interaction network.

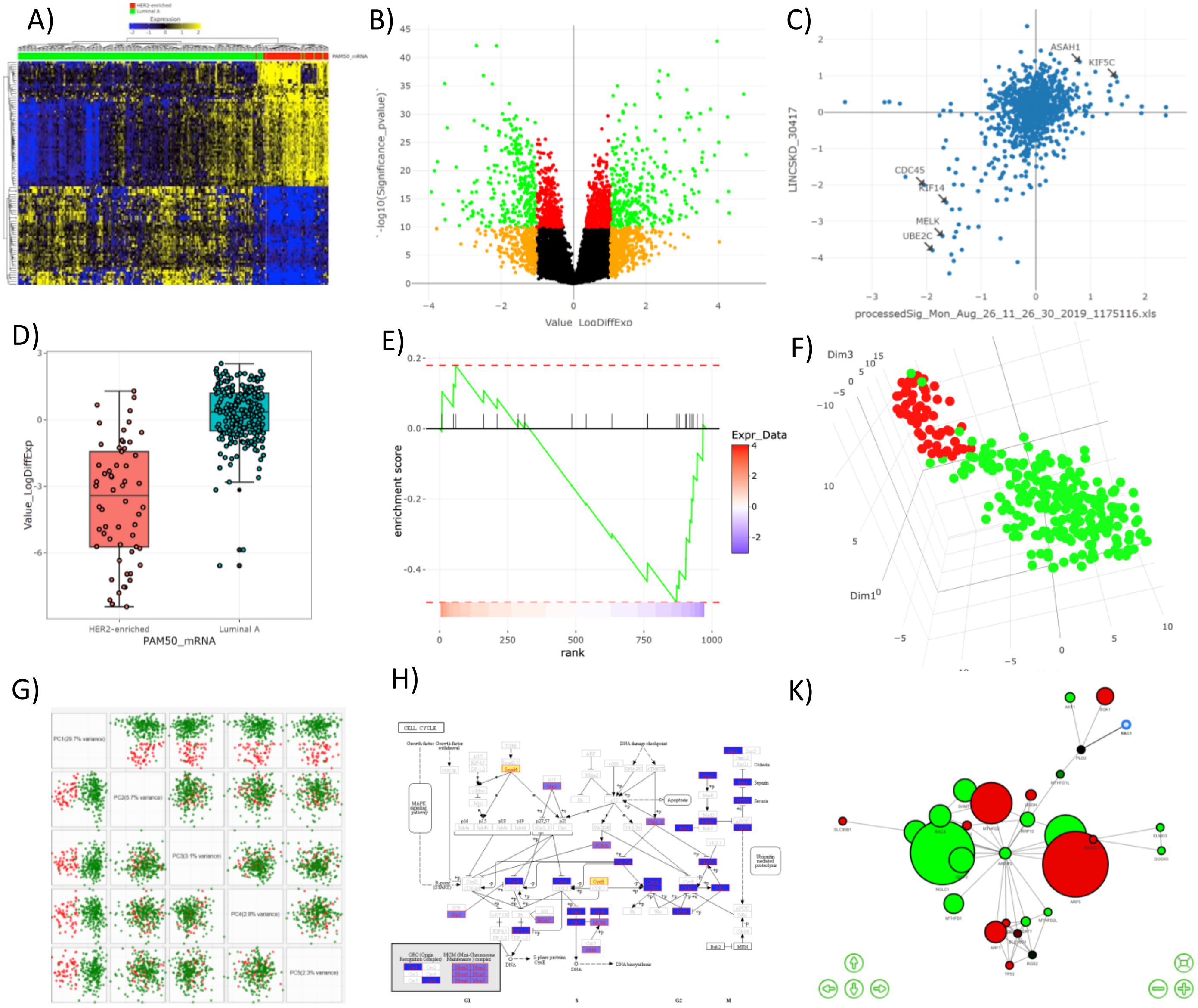

### Supplemental Figure 2: iLINCS architecture.

iLINCS is based on software stack: MySQL, MongoDB, R, NodeJS, and AngularJS. MySQL and MongoDB backend databases contain pre-processed genomics datasets, signatures and their connections, and all associated metadata. For external users, the database is accessed through a powerful API, which can be explored and tested with the Swagger UI interface. The iLINCS API was created in nodeJS using ExpressJS and Loopback frameworks. The API is documented and can be explored through swagger UI interface, which allows users to explore available API models and methods. The iLINCS internal analytical engine is written in R utilizing a range of specialized R packages. The iLINCS API is based on nodeJS using ExpressJS and Loopback frameworks. The nodeJS API connects with the R analytical engine using the openCPU framework which provides HTTP API for executing R functions. The user interfaces are written in Java Script using AngularJS framework. iLINCS facilitates submission of signatures and intermediate analysis results via redirection APIs to a range of third-party task specific bioinformatics web tools and services.

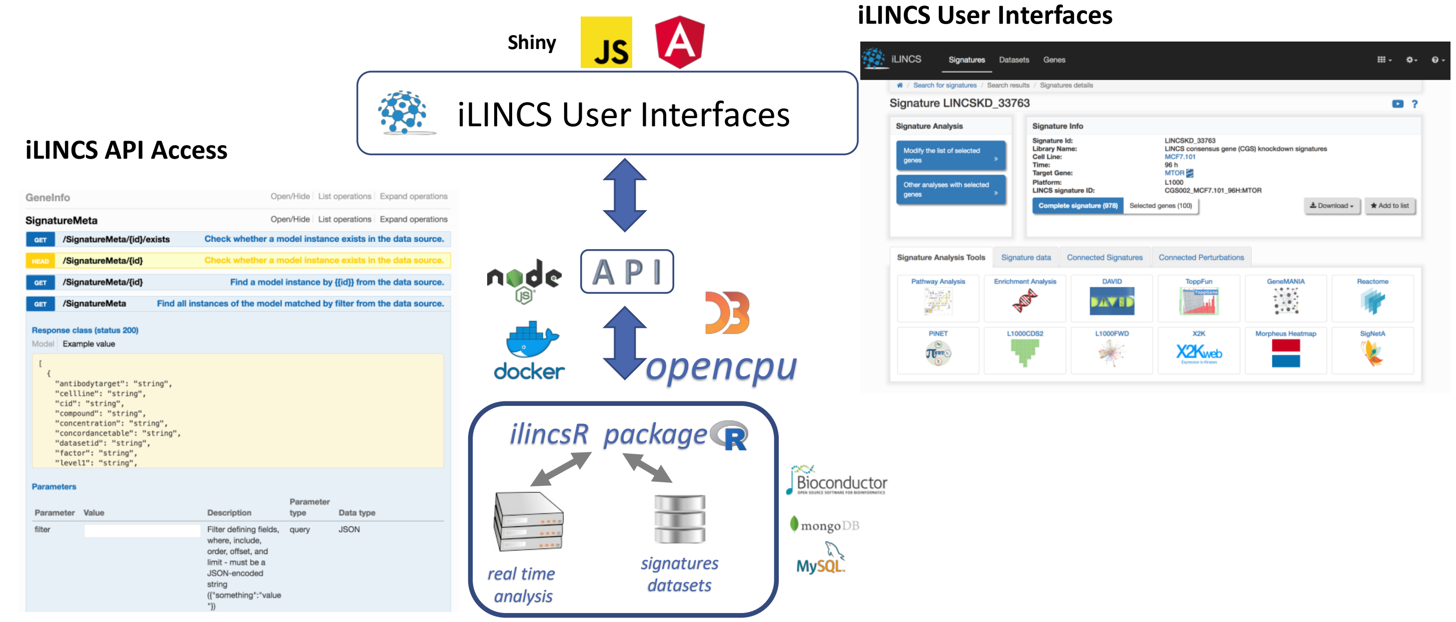

### Supplemental Results 1: Comparison of iLINCS and clue.io query results

iLINCS and clue.io query tools were compared based on the “connected” chemical perturbagens (CPs) returned by the query with the MTOR CRISPR consensus gene signature (CGS) used in the Use Case 1 of the main text (Fig 2A). For submission to the clue.io, the signature is summarized as the list of 50 most up-regulated and the 50 most down-regulated genes. The default clue.io connectivity metric, median τ across all cell-lines, was used to identify 20 most connected CPs and results, along with results obtained from iLINCS for the same CPs are shown in Table S1. Most of the connected CPs are also deemed significant by iLINCS analysis. Interestingly, none of the CPs implicated by clue.io, but not picked up as connected by iLINCS target the core elements of the mTOR signaling. On the other hand, all top 20 signatures (lowest pValue) implicated by iLINCS (Table S2) are known to target core elements of mTOR signaling (mTOR, PI3K and AKT proteins). This includes four perturbagens not deemed connected by clue.io (highlighted). Two CPs that show in the iLINCS results, but were missed by clue.io are not in the Touchstone set used by clue.io [4] (is_touchstone=0). Finally, 6 CPs are missing connectivity information in clue.io because they were part of the Phase II LINCS L1000 dataset (GSE70138), which are included in iLINCS, but not used by clue.io. The CPs in red were missing MOA information in the released data, and we established their MOA by manual literature and online databases searches.

In summary, clue.io and iLINCS provide qualitatively similar results with connected CPs targeting mTOR pathway. Some differences were introduced by the different statistical measure of similarity used by the two system, but the biggest differences most likely came from the scope of the query, where clue.io searches over a space of 2,837 Touchstone CPs and iLINCS searches over a space of 15,349 CPs with at least one high quality signature.

### Table S1: Top 20 CPs returned by the clue.io

| Perturbagen ID | Perturbagen Name | ilincsMoa | is_touchstone | median_tau | iLINCS | pValue |
|---|---|---|---|---|---|---|
| BRD-K67566344 | KU-0063794 | MTOR inhibitor | 1 | 97.35 | Connected | 3.83E-89 |
| BRD-K84937637 | sirolimus | MTOR inhibitor | 1 | 97.31 | Connected | 7.21E-55 |
| BRD-A84045418 | calpeptin | Calpain inhibitor | 1 | 97.23 | Connected | 2.08E-33 |
| BRD-K12184916 | NVP-BEZ235 | MTOR inhibitor | 1 | 96.02 | Connected | 6.94E-161 |
| BRD-K71726959 | BRD-K71726959 | CDK inhibitor | 1 | 95.33 | Not Connected | |
| BRD-K27305650 | LY-294002 | MTOR inhibitor | 1 | 95.32 | Connected | 3.35E-45 |
| BRD-K92577649 | GBR-13069 | Dopamine uptake inhibitor | 1 | 95.14 | Not Connected | |
| BRD-K30677119 | PP-30 | RAF inhibitor | 1 | 94.71 | Connected | 3.48E-18 |
| BRD-K99818283 | PIK-90 | PI3K inhibitor | 1 | 94.42 | Connected | 1.27E-54 |
| BRD-K69932463 | AZD-8055 | MTOR inhibitor | 1 | 93.54 | Connected | 1.17E-218 |
| BRD-A62025033 | temsirolimus | MTOR inhibitor | 1 | 93.49 | Connected | 3.46E-125 |
| BRD-K77008974 | WYE-354 | MTOR inhibitor | 1 | 92.05 | Connected | 4.03E-19 |
| BRD-K06593056 | LE-135 | Retinoid receptor agonist | 1 | 91.85 | Not Connected | |
| BRD-K21350491 | phenamil | TRPV antagonist | 1 | 91.54 | Not Connected | |
| BRD-K64835161 | BRD-K64835161 | CLK inhibitor,DYRK inhibitor | 1 | 91.53 | Connected | 1.18E-04 |
| BRD-A77299732 | salubrinal | Eukaryotic translation initiation factor inhi | 1 | 91.47 | Not Connected | |
| BRD-K13800121 | parecoxib | Cyclooxygenase inhibitor | 1 | 90.63 | Connected | 5.08E-06 |
| BRD-K67868012 | PI-103 | PI3K inhibitor,MTOR inhibitor | 1 | 90.59 | Connected | 7.37E-144 |
| BRD-K71879491 | tretinoin | Retinoid receptor agonist | 1 | 90.33 | Not Connected | |
| BRD-A75409952 | wortmannin | PI3K inhibitor | 1 | 90.30 | Connected | 2.14E-110 |

### Table S2: Top 20 CPs returned by the iLINCS

| Perturbagen ID | Perturbagen Name | ilincsMoa | is_touchstone | median_tau | iLINCS | pValue |
|---|---|---|---|---|---|---|
| BRD-K02708799 | GSK 1059615 | PI3K inhibitor | 0 | | Connected | 0 |
| BRD-A79768653 | sirolimus | MTOR inhibitor | 1 | 83.07 | Connected | 1.6E-293 |
| BRD-K69932463 | AZD-8055 | MTOR inhibitor | 1 | 93.54 | Connected | 1.2E-218 |
| BRD-K12184916 | NVP-BEZ235 | MTOR inhibitor | 1 | 96.02 | Connected | 6.9E-161 |
| BRD-K59317601 | MLN-0128 | MTOR inhibitor | | | Connected | 4.6E-144 |
| BRD-K67868012 | PI-103 | PI3K inhibitor,MTOR inhibitor | 1 | 90.59 | Connected | 7.4E-144 |
| BRD-K94012289 | 936890-98-1 | MTOR inhibitor | 1 | | Connected | 9.4E-137 |
| BRD-K40175214 | torin-1 | MTOR inhibitor,PI3K inhibitor | 1 | 71.90 | Connected | 1.9E-135 |
| BRD-A45498368 | WYE-125132 | MTOR inhibitor | 1 | 79.43 | Connected | 4.3E-127 |
| BRD-A62025033 | temsirolimus | MTOR inhibitor | 1 | 93.49 | Connected | 3.5E-125 |
| BRD-K52911425 | GDC-0941 | PI3K inhibitor | 1 | 82.75 | Connected | 5.2E-125 |
| BRD-A75409952 | wortmannin | PI3K inhibitor | 1 | 90.30 | Connected | 2.1E-110 |
| BRD-K13154216 | Everolimus | MTOR inhibitor | | | Connected | 4.4E-109 |
| BRD-K42898655 | Temsirolimus | MTOR inhibitor | | | Connected | 8E-104 |
| BRD-K99023089 | AZD5363 | AKT inhibitor | | | Connected | 3.2E-100 |
| BRD-K09078998 | Ridaforolimus | MTOR inhibitor | | | Connected | 8.83E-97 |
| BRD-K63068307 | ZSTK-474 | PI3K inhibitor | 1 | 78.95 | Connected | 7.61E-94 |
| BRD-K67566344 | KU-0063794 | MTOR inhibitor | 1 | 97.35 | Connected | 3.83E-89 |
| BRD-K72636697 | SCHEMBL17052537 | MTOR inhibitor | 0 | | Connected | 2.91E-86 |
| BRD-A25736793 | Everolimus | MTOR inhibitor | | | Connected | 4.02E-79 |

### Supplemental Results 2: Benchmarking methods for connectivity analysis and comparisons to other web resources

We performed a limited benchmarking study of several methods for "connecting" transcriptional signatures with the objective to identify optimal methods for pre-computed connections between signatures in the iLINCS libraries, and for connectivity analysis of newly created and submitted signatures. The choice of the methods were motivated by previous studies[4,6-8] and included Extreme and Weighted correlations, the Connectivity Score used by the Connectivity Map, and the correlation based on signed log significance, which was motivated by our previous results in the context of enrichment analysis[10]. The methods were benchmarked with respect to their ability to "connect" a chemical perturbagen signature in one cancer cell line with the equivalent signature in another cell line for five cell lines with most signatures in the LINCS L1000 dataset (A375, A549, PC3, VCAP and MCF7). Only signatures designated as "gold" and generated using the "epsilon" version of L1000 probes (GEO CMap LINCS User Guide v2.1) were used in the analysis. Two signatures that share the same treatment (same small molecule, same dose and duration) in two different cell lines were considered to be true positive signature pair. True negative signature pairs were defined as pairs of signatures from different cell lines treated with different chemical perturbagen. For each 10 different cell line combinations, the number of "true positive pairs" ranged between 5,793 and 16,477 (the average number is 10,145). To calculate ROC curves, 50,000 "true negative pairs" randomly selected from all true negative pairs for a given cell line combination. Following six methods were compared:

**WeightedCor:** Weighted Pearson's correlation between MODZ scores, weighted by the logarithm of the product of the p-values. This score was shown to be best in correlating genome wide signatures of related biological states [8].

**Cor:** Pearsons's correlation between MODZ scores of the two signatures.

**ExtremeCorLobP_100**: Extreme Correlation of Signed Log P-values utilizing top 100 up- and down-regulated genes. The signed significance of the i[th] gene is defined as

$$s_i = sign(d_i) * (-log_{10}(p_i)),\ \ for\ \ i = 1, \dots, N,$$

and the signed significance signature is $\boldsymbol{s}=(s_1,\dots,s_N)$. The extreme signed signature $\boldsymbol{e}=(e_1,\dots,e_N)$ is then constructing by setting the signed significances of all genes other than the top 100 and bottom 100 to zero:

$$e_i = \begin{Bmatrix} s_i, if\ s_i \geq s^{100}\ or\ s_i \leq s^{-100} \\ 0, otherwise \end{Bmatrix}$$

Where $s^{100}$ is the 100th most positive $s_i$ and $s^{-100}$ is the 100th most negative $s_i$. The extreme Pearson correlation between two signatures is then calculated as the standard Pearson's correlation between the extreme signed significance signatures.

**ExtremeWeightedCor_100**: Extreme Weighted Pearson's Correlations calculating weighted Pearson's correlation (see **WeightedCor**) after setting MODZ's of all but top 100 up- and down-regulated genes to zero (see **ExtremeCorLobP_100)**

**CMap_100:** Connectivity score as described in the Connectivity Map publication [4] using the top 100 up- and down-regulated genes as the query.

Overall, all six methods performed very well on this task with average AUC's ranging from 0.946 to 0.926 (Table S3). The decision was made to use the Extreme Correlation of Signed Log P-values utilizing top 100 up- and down-regulated genes as the method for pre-computed connections, since it showed the best performance. For real-time connectivity analysis of submitted and newly created signatures, we decided to use Weighted Correlations since they were almost as good as the best method, but much easier to compute in the real time. To facilitate the fast computations of weighted correlations, the algorithm was implemented in C and added to the backend R package using the *Rcpp* package [13].

### Table S3: Area under the ROC curve (AUC) for six competing connectivity analysis methods across 10 pairs of cancer cell lines.

| Method | A375xA549 | A375xMCF7 | A375xPC3 | A375xVCAP | A549xMCF7 | A549xPC3 | A549xVCAP | MCF7xPC3 | MCF7xVCAP | PC3xVCAP | Average |
|---|---|---|---|---|---|---|---|---|---|---|---|
| ExtremeCorLogP_100 | 0.951 | 0.952 | 0.957 | 0.930 | 0.947 | 0.947 | 0.926 | 0.957 | 0.947 | 0.948 | 0.946 |
| ExtremeWeightedCor_100 | 0.946 | 0.947 | 0.951 | 0.925 | 0.944 | 0.944 | 0.921 | 0.953 | 0.944 | 0.946 | 0.942 |
| WeightedCor | 0.945 | 0.947 | 0.952 | 0.924 | 0.943 | 0.943 | 0.918 | 0.954 | 0.941 | 0.943 | 0.941 |
| ExtremeCor_100 | 0.942 | 0.946 | 0.947 | 0.921 | 0.941 | 0.941 | 0.914 | 0.952 | 0.936 | 0.939 | 0.938 |
| Cor | 0.938 | 0.943 | 0.945 | 0.914 | 0.936 | 0.937 | 0.903 | 0.949 | 0.925 | 0.929 | 0.932 |
| CMap_100 | 0.935 | 0.940 | 0.941 | 0.906 | 0.931 | 0.933 | 0.895 | 0.945 | 0.915 | 0.921 | 0.926 |

### Table S4: Comparing the scope and functionality of iLINCS with other resources.

There are numerous online analysis tools that cover different aspects of cancerLINCS functionality: access to a vast amount of data[2,9,11,14,15], "connectivity analysis" with LINCS signatures[4,9], user friendly web interfaces for analysis of primary transcriptomic data[16], and more targeted collections of cancers omics datasets[1,3,5], various aspects of systems biology analysis of omics signatures[17-23], and interactive visualizations[24,25]. However, *iLINCS* platform is unique in bringing together all different aspects of a comprehensive signature analysis platform.

| | *GUI based analysis* | *API* | *Signature Creation* | *Systems Biology Analysis* | *Connectivity Analysis* | *Multi-omic Analysis* | *Bulk Transcriptomic Datasets* | *Single Cell RNA-seq Datasets* | *Proteomic Datasets* |
|---|---|---|---|---|---|---|---|---|---|
| *iLINCS* | **Yes** | **Yes** | **Yes** | **Yes** | **Yes** | **Yes** | **Yes** | **Limited** | **Yes** |
| *cBioPortal*[1] | **Yes** | **Yes** | **No** | **No** | **No** | **No** | **Yes** | **No** | **No** |
| *GDC*[2] | **Yes** | **Yes** | **No** | **No** | **No** | **No** | **Yes** | **No** | **No** |
| *Expression Atlas*[3] | **Limited** | **Limited** | **Limited** | **No** | **No** | **No** | **Yes** | **Limited** | **No** |
| *CancerSEA*[5] | **Yes** | **No** | **No** | **No** | **No** | **No** | **No** | **Limited** | **No** |
| *BioJupies*[9] | **No** | **No** | **Yes** | **Yes** | **Yes** | **No** | **Yes** | **No** | **No** |
| *GEO*[11] | **Yes** | **Limited** | **Limited** | **No** | **No** | **No** | **Yes** | **Limited** | **Limited** |
| *clue.io*[4] | **Yes** | **Limited** | **No** | **No** | **Yes** | **No** | **No** | **No** | **No** |
| *WebMeV*[12] | **Yes** | **No** | **Yes** | **Yes** | **No** | **No** | **Yes** | **No** | **No** |
